# Perturbation of ribosomal subunits dynamics by inhibitors of tRNA translocation

**DOI:** 10.1101/2021.03.23.436553

**Authors:** Riccardo Belardinelli, Heena Sharma, Frank Peske, Marina V. Rodnina

## Abstract

Many antibiotics that bind to the ribosome inhibit translation by blocking the movement of tRNAs and mRNA or interfering with ribosome dynamics, which impairs the formation of essential translocation intermediates. Here we show how translocation inhibitors viomycin (Vio), neomycin (Neo), paromomycin (Par), kanamycin (Kan), spectinomycin (Spc), hygromycin B (HygB), and streptomycin (Str, an antibiotic that does not inhibit tRNA translocation), affect principal motions of the small ribosomal subunits (SSU) during EF-G-promoted translocation. Using ensemble kinetics, we studied the SSU body domain rotation and SSU head domain swiveling in real time. We show that although antibiotics binding to the ribosome can favor a particular ribosome conformation in the absence of EF-G, EF-G-induced transition to the rotated/swiveled state of the SSU is hardly affected. The major effect of the antibiotics is observed at the stage when the SSU body and the head domain move backward. Vio, Spc and high concentrations of Neo completely inhibit the backward movements of the SSU body and head domain. Kan, Par, HygB and low concentrations of Neo slow down both movements, but their sequence and coordination are retained. Finally, Str has very little effect on the backward rotation of the SSU body domain, but retards the SSU head movement. The data underscore the importance of ribosome dynamics for tRNA-mRNA translocation and provide new insights into the mechanism of antibiotic action.

## Introduction

The bacterial ribosome is a major target for antibiotic inhibitors of protein synthesis. Ribosome-targeting antibiotics mostly bind to the functional centers of the two ribosomal subunits, such as the decoding center, the peptidyl transferase center, or the polypeptide exit tunnel, and obstruct the ribosome function in different ways (Yonath 2005; Wilson 2009). As many antibiotics have several binding sites, they can simultaneously inhibit several steps of translation, leading to pleiotropic concentration-dependent effects. One of the most common mechanisms of antibiotic action is to block the essential dynamics of the ribosome. Among the four steps that encompass translation, i.e. initiation, elongation, termination, and ribosome recycling, translocation is arguably the most dynamic (Belardinelli et al. 2016b; Noller et al. 2017a). In each round of elongation, ribosomes cycle between two main conformational states that differ in the relative orientation of the ribosomal subunits designated as the non-rotated (N) state and rotated (R) state (Chen et al. 2012; Noller et al. 2017b; Rodnina et al. 2019). Upon transition from the N to the R state, the body domain of the small ribosomal subunit (SSU) moves relative to the large ribosomal subunit (LSU) in a counterclockwise (forward) direction when looking from the SSU. At the same time, the SSU head domain rotates counterclockwise relative to the SSU body to a swiveled (S) state. As the ribosomal subunits rotate, the tRNAs move from the classical (C) to hybrid (H) states that differ in the positions of the tRNA 3’ ends on the LSU. The ribosome with a peptidyl-tRNA in the P site favors the N/C state. Binding of aminoacyl-tRNA to the A site of the ribosome and the incorporation of the incoming amino acid into the polypeptide nascent chain yields a ribosome complex with a deacylated tRNA in the P site and a peptidyl-tRNA in the A site, which is ready for the subsequent translocation and is called a pretranslocation (PRE) complex. Ribosomes in the PRE complex favor the R/H state, although the distribution between the N and R states depends on the tRNA bound to the ribosome and experimental conditions (Sharma et al. 2016).

Translocation is facilitated by the ribosomal translocase elongation factor G (EF-G) (Figure 1A). Binding of EF-G to the PRE complex accelerates the forward movement and stabilizes the complexes in R state (Sharma et al. 2016). In the following translocation reaction, the tRNAs move from the A to the P and from the P to the E site, from which the deacylated tRNA is released. This process requires a major backward movement of the subunits from the R to the N state and proceeds in several steps (Belardinelli et al. 2016b; Noller et al. 2017a). Importantly, the timing of the SSU body and head domain movement is uncoupled (Belardinelli et al. 2016a). The SSU body rotates from the R back towards the N state at an early step of translocation, whereas the reverse movement of the SSU head domain (i.e. S to N, this time non-swiveled) is delayed (Belardinelli et al. 2016a). Uncoupling of these motions leads to the formation of a key translocation intermediate in which the tRNAs move towards the P and the E site (Ramrath et al. 2013; Zhou et al. 2014; Belardinelli et al. 2016a; Peng et al. 2019). Further backward movement of the SSU head domain into the ground state completes translocation and coincides with the release of EF-G and the E-site tRNA from the ribosome and results in the formation of the posttranslocation (POST) state (Belardinelli et al. 2016a).

**Figure 1.**
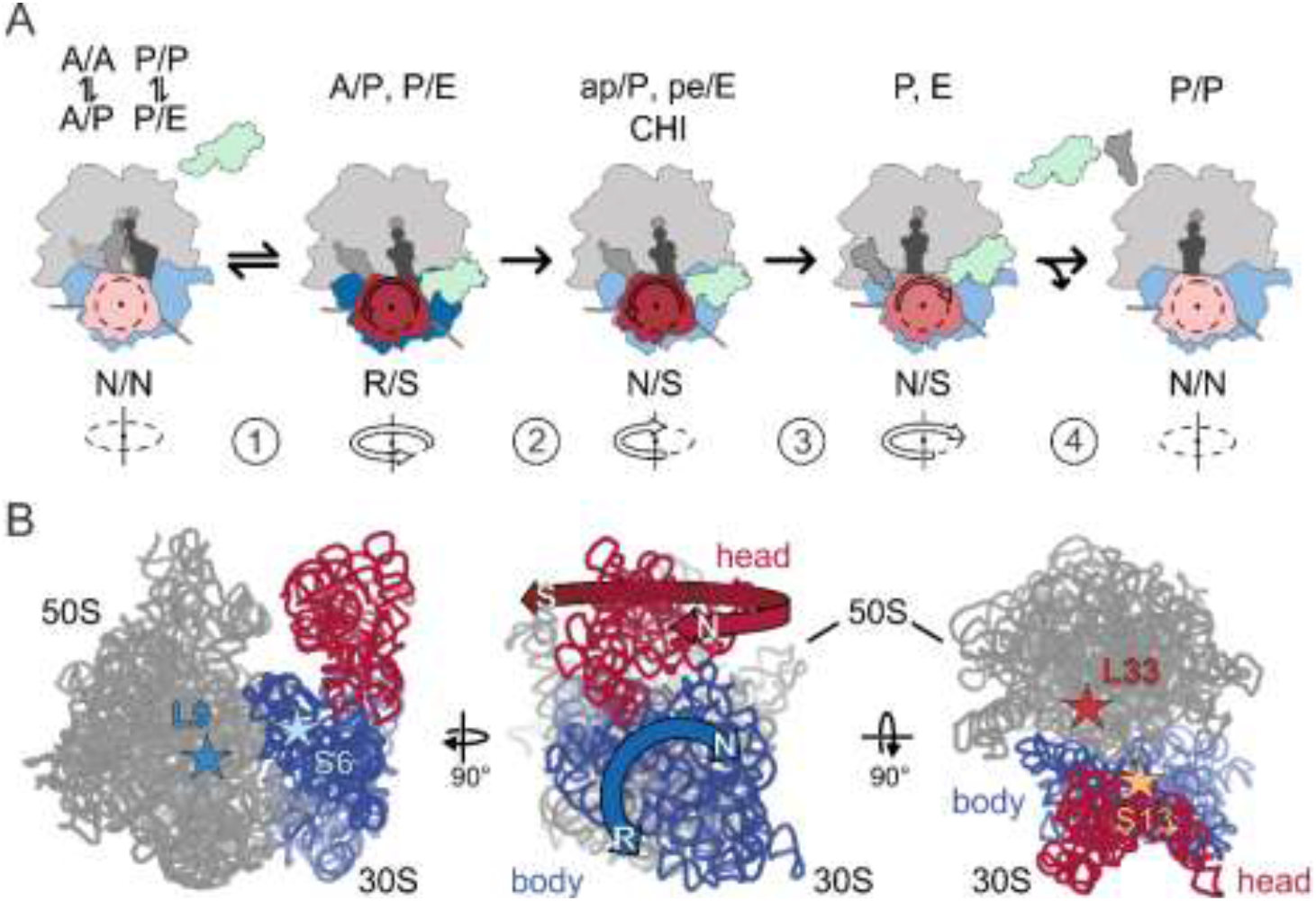
Schematic of SSU body rotation and head swiveling. (A) Simplified schematic of tRNA-mRNA translocation. (1) EF-G binding stabilizes the R/S state of the ribosome; (2) Pi release facilitates the backward movement of the SSU body domain towards the N state, whereas the head domain remains in the S state; the tRNAs move to the CHI state. (3) The SSU head domain starts to move backwards and tRNAs translocate to the P and E sites. (4) The ribosome adopts an N state and EF-G and the E-site tRNA dissociate. (B) Left panel, positions of the S6-L9 FRET pair that reports on the SSU body rotation (blue) relative to the LSU (gray). Center panel, directions of the principal movements of the SSU body and head domains. Right panel, positions of the S13-L33 FRET pair to monitor the swiveling of the SSU head (red) relative to the body. Ribosomal proteins are omitted for graphical simplicity. Stars indicate the position of the fluorescence labels on the ribosomal proteins.

The most common antibiotics inhibiting translation are viomycin (Vio), neomycin (Neo), paromomycin (Par), kanamycin (Kan), spectinomycin (Spc), hygromycin B (HygB), streptomycin (Str), and fusidic acid (FA). Vio is a cyclic peptide that has several distinct binding sites on the ribosome (Zhang et al. 2020). Two Vio molecules bind to the SSU near the decoding site (Stanley et al. 2010; Zhang et al. 2020). The other three Vio molecules bind at the interface between the SSU and LSU and stabilize the R state (Zhang et al. 2020). Neo, Par, Kan, Spc, and HygB are aminoglycosides that bind near the decoding site of the ribosome with details of binding depending on the antibiotic. Par binds the SSU helix 44 of 16S rRNA (Vicens and Westhof 2001; Selmer et al. 2006; Watson et al. 2020) and stabilizes the N state. According to recent structural data, it also interacts with LSU Helix 69 of 23S rRNA (Wang et al. 2012; Wasserman et al. 2015). Similar to Par, Neo can bind to both subunits at helices SSU h44 and LSU H69, respectively, however, in contrast to Par, Neo stabilizes an intermediate state of subunits rotation (Wasserman et al. 2015). Kan belongs to the same class as Par and Neo and binds the decoding center in the same binding pocket formed by the nucleotides 1492-1493 of 16S rRNA. FA is a steroid antibiotic, which – in contrast to other drugs inhibiting translocation– binds to EF-G, rather than to the ribosome, and blocks EF-G dissociation from the POST complex (Wasserman et al. 2016). Because its mechanism of action is well studied (Spiegel et al. 2007; Borg et al. 2015; Belardinelli and Rodnina 2017) and is distinct from that of other antibiotics blocking translocation, we do not discuss the FA action in the following.

Although all of these antibiotics inhibit translocation, they act by somewhat different mechanisms. In principle, antibiotic binding can stabilize a ribosome conformation that is unfavorable for EF-G binding (Wang et al. 2012). Antibiotics can also selectively stabilize the N or R state, thereby inhibiting dynamics of the ribosome during translocation, or increase the affinity of tRNA binding to the A site, thereby preventing the tRNA from moving into the P site (Peske et al. 2004). The existence of the secondary binding sites may complicate the picture even further. For example, the effect of Neo on subunit rotation depends on the antibiotic concentration in a non-linear biphasic way, suggesting the role of secondary binding sites at high antibiotic concentration (Feldman et al. 2010). The antibiotics may also inhibit the swiveling motions of the SSU head domain, but this has not been studied so far. Finally, the effect of antibiotics on the kinetics of SSU movements is not known. The goal of the present study is to systematically test the effects of translocation inhibitors on the real-time movements of the SSU body and head domains and to combine these results into a coherent picture of antibiotic action in EF-G-dependent translocation.

## Results

### Experimental approach: ensemble kinetics

We studied the effect of the antibiotics upon EF-G–GTP-facilitated translocation by ensemble kinetics experiments using a stopped-flow technique. Previous ensemble experiments demonstrated that after EF-G binding, GTP hydrolysis and N to R transition are very rapid (250 s^−1^ and 200 s^−1^, respectively, at saturating EF-G concentrations) (Savelsbergh et al. 2003; Sharma et al. 2016). Here we monitor the steps of SSU body and head domain rotation (Belardinelli et al. 2016a) (Figure 1B). As antibiotics dramatically alter ribosome dynamics, we used qualitative comparisons between the apparent rate constants of the SSU body rotation and SSU head swivel in the absence and presence of antibiotics, rather than deriving rate constants of each reaction step, as described in our previous reports (Belardinelli et al. 2016a; Sharma et al. 2016).

We monitored the SSU body rotation and SSU head swiveling by ensemble FRET measurements using two well-established FRET pairs, S6–L9 and S13–L33, respectively (Ermolenko et al. 2007a; Belardinelli et al. 2016a; Sharma et al. 2016). We prepared ribosomal subunits from deletion strains lacking one of these ribosomal proteins. To introduce fluorescence labels on the ribosomal subunits, the respective protein bearing a single cysteine was overexpressed, fluorescence labeled, and use to reconstitute the respective ribosomal subunit lacking this protein. Double-labeled S6–L9 or S13–L33 ribosomes were then used to prepare PRE complexes harboring deacylated tRNA^fMet^ in the P site and peptidyl-tRNA (fMet-Phe-tRNA^Phe^) in the A site. We used fluorophores Alexa568 and Alexa488 to label the S6–L9 pair and Atto540Q and Alexa488 to label the S13–L33 pair; for both pairs, the N state have higher FRET efficiency than the R and the S states (Belardinelli et al. 2016a; Sharma et al. 2016). Fluorescence labeling of ribosomal subunits did not impaired ribosome activity (Belardinelli et al. 2016a; Sharma et al. 2016).

In the absence of antibiotics, the addition of EF-G–GTP to the PRE complex results in a rapid forward rotation/swiveling of the SSU body and head domains (Figure 2B, blue trace; Table 1), followed by GTP hydrolysis by EF-G to GDP and inorganic phosphate, Pi. The SSU body domain rotates backward towards the N state, which is then followed by a slower reverse swiveling of the SSU head domain. Both reverse reactions appear biphasic (Figure 2B, blue trace: *k*_app2_ = 48 s^−1^ and *k*_app3_ = 8 s^−1^; red trace: *k*_app2_ = 8 s^−1^ and *k*_app3_ = 1 s^−1^) (Table 1). In an earlier study, we have shown that such apparently biphasic time courses represent a quasi-continuous movement of the SSU body and head (Belardinelli et al. 2016a). Here, for comparison between the curves, we calculated the weighted average lifetime τ, which in the absence of antibiotics is about 30 ms for the R to N movements of the SSU body and 190 ms for the backward movement of the SSU head (S to N) (Table 1). The SSU body reverse rotation coincides with the Pi release from EF-G and the movement of the tRNAs into the intermediate state of translocation, denoted as chimeric (CHI) (Savelsbergh et al. 2003; Zhou et al. 2014; Belardinelli et al. 2016a). In this complex, the A-site tRNA anticodon remains attached to the A-site elements of the SSU head domain, but changes the interactions on the SSU body domain due to its back rotation. The 3’ end of the A-site tRNA is in the P site on the LSU and the overall orientation of the A-site tRNA is denoted as ap/P. By analogy, the P-site tRNA moves into a pe/E state. The lifetime of the CHI state can be estimated from the difference in the time courses of backward SSU body rotation and head swiveling, τ_CHI_ = 162 ms (**Table 1**). Back swiveling of the SSU head domain largely coincides with the dissociation of the E-site tRNA and EF-G from the ribosome and represents late steps of translocation (Belardinelli et al. 2016a). This simplified kinetic model of translocation allows us to combine the present and previous results obtained with different reports to identify the detailed mechanism of antibiotic action on translocation. In the following section, we will describe these findings for each antibiotic or antibiotics group.

**Table 1.** Apparent rate constants of SSU body and head rotation in the presence of EF-G and different antibiotics. k_app1_ represents the forward rotation/swivel; k_app2_ and k_app3_ represent the backward reaction towards the N state.

| Antibiotic | Body rotation | | | | Head swiveling | | | | $\tau_{CHI}$<br>(s) |
| --- | --- | --- | --- | --- | --- | --- | --- | --- | --- |
| | $\tau_{body}$<br>(s) | $k_{app1}$<br>(s<sup>-1</sup>) | $k_{app2}$<br>(s<sup>-1</sup>) | $k_{app3}$<br>(s<sup>-1</sup>) | $\tau_{head}$<br>(s) | $k_{app1}$<br>(s<sup>-1</sup>) | $k_{app2}$<br>(s<sup>-1</sup>) | $k_{app3}$<br>(s<sup>-1</sup>) | |
| **None** | 0.029 ± 0.004 | 699 ± 153 | 47.1 ± 2.4 | 8.2 ± 0.7 | 0.191 ± 0.008 | 159 ± 7 | 7.6 ± 0.2 | 1.2 ± 0.1 | 0.162 ± 0.012 |
| **Spc** | 10.9 ± 2.3 | 34 ± 41 | 0.34 ± 0.05 | 0.033 ± 0.001 | 46 ± 2 | 125 ± 28 | 0.09 ± 0.01 | 0.009 ± 0.001 | 35 ± 8 |
| **Par** | 0.56 ± 0.01 | 105 ± 7 | 1.8 ± 0.1 | n.a. | 2.6 ± 0.1 | 56 ± 3 | 0.53 ± 0.01 | 0.054 ± 0.001 | 2.1 ± 0.1 |
| **Kan** | 0.83 ± 0.48 | 42 ± 2 | 1.4 ± 0.1 | 0.47 ± 0.19 | 4.9 ± 0.3 | 39 ± 1 | 0.29 ± 0.01 | 0.050 ± 0.002 | 4.1 ± 0.8 |
| **HygB** | 17.6 ± 7.6 | 36 ± 14 | 0.12 ± 0.02 | 0.034 ± 0.002 | 153 ± 66 | 20 ± 4 | 0.016 ± 0.002 | 0.004 ± 0.001 | 136 ± 73 |
| **Neo<sub>0.2</sub>** | 1.45 ± 0.03 | 95 ± 8 | 0.69 ± 0.01 | n.a. | 4.5 ± 0.2 | 53 ± 2 | 0.31 ± 0.01 | 0.046 ± 0.001 | 3.1 ± 0.2 |
| **Str** | 0.07 ± 0.03 | 86 ± 14 | 17 ± 3 | 5.1 ± 1.4 | 46 ± 6 | 124 ± 28 | 0.09 ± 0.01 | 0.009 ± 0.001 | 46.2 ± 6 |

**Figure 2.**
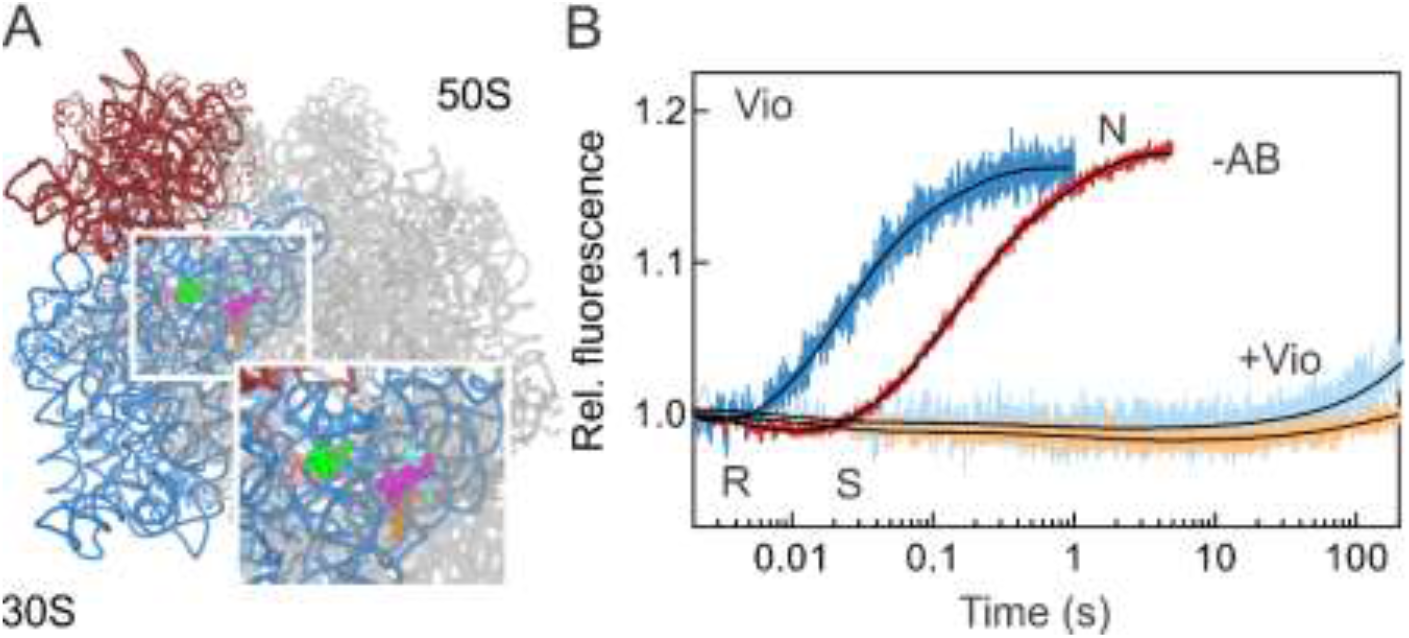
Vio inhibits formation of CHI state and all following events of translocation. (A) Vio binding sites on the SSU (pink and light green) and LSU (orange, magenta and cyan) (Zhang et al. 2020). SSU body and head are highlighted in blue and red, respectively; LSU is gray. (B) EF-G-dependent SSU body rotation and head swiveling were monitored in the absence of antibiotic (−AB; blue and red, respectively) or in the presence of Vio (light blue and orange, respectively). Conformations of the SSU body and head domains (N, R and S) are indicated. Here and in all subsequent experiments, PRE complexes assembled with double-labeled ribosomes (0.05 μM after mixing) were mixed with EF-G (4 μM) and GTP (0.5 mM).

### Viomycin

Vio is a strong inhibitor of tRNA–mRNA translocation (Liou and Tanaka 1976; Modolell and Vazquez 1977; Peske et al. 2004; Ermolenko et al. 2007b; Bulkley et al. 2012; Wilson 2014; Zhang et al. 2020). Vio binding to the PRE complex dramatically increases the affinity of peptidyl-tRNA binding to the A site (Peske et al. 2004), most probably due to the Vio molecule bound to the decoding site at the bridge b2a (Zhang et al. 2020). The equilibrium between the N and R conformations shifts towards the R state with the SSU head domain in a swiveled conformation (Wang et al. 2012). The rotation is facilitated by the three Vio molecules bound at the ribosome interface (Zhang et al. 2020). The fraction of tRNAs in the hybrid state also increases (Cornish et al. 2008).

Vio does not abolish EF-G binding to the ribosome and in fact seems to stabilize EF-G binding (Salsi et al. 2014), although the association rate appears slower than in the absence of the antibiotic (Belardinelli et al. 2016a). GTP hydrolysis by EF-G and Pi release are not affected (Savelsbergh et al. 2003) and the 3’ end of the A-site tRNA moves towards the P site on the LSU (Holtkamp et al. 2014). However, the backward movement of the SSU body and head domains are completely blocked (Figure 2A, light blue and light red curves). Thus, Vio blocks translocation by stabilizing the A-site tRNA and by interfering with the back rotation of the ribosomal subunits. Notably, multiple turnover GTPase activity of EF-G is only slightly affected (Peske et al. 2004), indicating that Vio does not block EF-G on the ribosome.

### Spectinomycin

Crystal structures show Spc binding to h34 near the neck helix of the SSU head domain (Carter et al. 2000; Borovinskaya et al. 2007b) (Figure 3A). Unlike Vio, Spc destabilizes the A-site tRNA binding, which *per se* should be favorable for translocation (Peske et al. 2004). In the absence of EF-G, Spc impedes the dynamics of the SSU head swiveling and stabilizes the non-swiveled state (Borovinskaya et al. 2007b).

**Figure 3.**
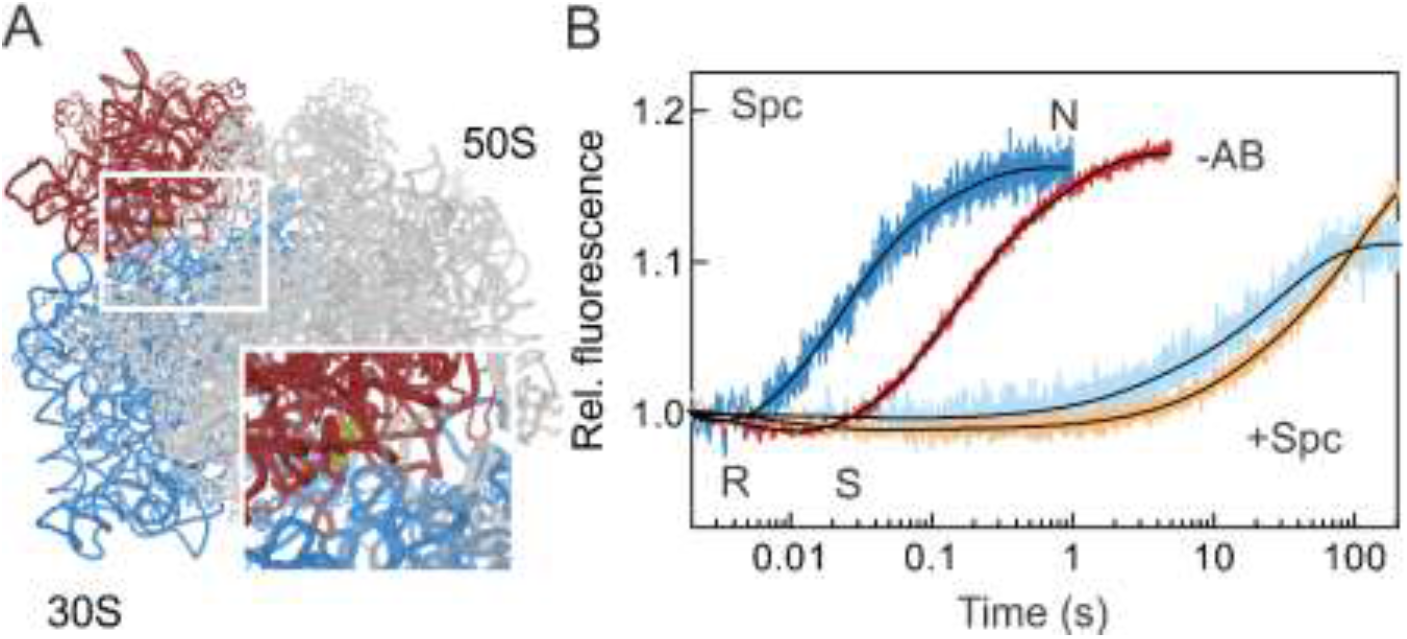
Spc action on ribosome dynamics. (A) Spc binding site on the SSU is indicated in olive. SSU body and head are highlighted in blue and red, respectively, LSU in gray. (B) EF-G dependent body (blue shade) and head rotation dynamics (orange shade) were monitored in the presence of Spc (lighter shades), and compared to the reaction in the absence of the drug (-AB; darker shades). Conformations of the SSU body and head domains (N, R and S) are indicated.

Spc does not affect EF-G binding to the ribosome, GTP hydrolysis, Pi release, or the movement of the 3’ end of the A-site peptidyl-tRNA towards the P site (Peske et al. 2004; Holtkamp et al. 2014). It does not inhibit the movement of the P-site tRNA towards the E site, leading to the formation of the so-called INT state (Pan et al. 2007). However, the backward rotation of the SSU body is inhibited (Figure 3B), consistent with single molecule FRET measurements (Aitken and Puglisi 2010; Chen et al. 2012; Wasserman et al. 2015; Wasserman et al. 2016). Backward movement of the SSU body (τ = 11 s) becomes the rate-limiting step of translocation in the presence of Spc, although the back swiveling of the SSU head domain is also very slow τ = 46 s) (Fig. 3B and Table 1). Spc allows slow tRNA–mRNA translocation (Peske et al. 2004). Ensemble kinetic measurements suggest that in the presence of Spc, slow translocation proceeds via a pathway that is also observed in the absence of Spc on a small fraction of the ribosomes. As the Spc concentration increases, a larger fraction of ribosomes changes to the slow regime (Peske et al. 2004). Interestingly, a similar slow regime, with a slow back rotation or the SSU body, has been observed in the absence of antibiotics when GTP was replaced by a non-hydrolysable analog GDPNP (Belardinelli et al. 2016a). Thus, it is tempting to speculate that Spc, similarly to non-hydrolysable GTP analogs, uncouples the conformational changes induced upon GTP hydrolysis and Pi release from the back rotation of the SSU body. The resulting slow translocation proceeds through a different pathway, which would also explain the observation from smFRET experiments, suggesting that the CHI state is not sampled in the presence of Spc (Wasserman et al. 2015).

### Kanamycin, paromomycin, and hygromycin B

Kan, Par, and HygB are aminoglycoside antibiotics that bind near the decoding center. Kan and Par are 4,5-and 4,6-aminoglycosides, respectively. Kan binds to 16S rRNA at nucleotides 1491 and 1408, whereas Par has two known binding sites, one on the SSU in h44 of 16S rRNA (Selmer et al. 2006; Watson et al. 2020) and one in the LSU 23S rRNA (Feldman et al. 2010; Wasserman et al. 2015). Binding of Kan or Par to PRE complexes at the SSU sites stabilizes the N state (Selmer et al. 2006; Feldman et al. 2010; Wang et al. 2012; Watson et al. 2020). Additionally, when Par binds to both binding sites on the SSU and the LSU, the P-site tRNA moves to the P/pe state and the ribosome adopts an intermediate state of rotation (Tsai et al. 2013; Wasserman et al. 2015). HygB binds at the apical part of helix 44 and contacts nucleotides 1493, 1494 and 1405 (Borovinskaya et al. 2008). Two conformations are observed in crystal structures of vacant ribosomes in the presence of HygB. Both structures present the body in a non-rotated conformation (−2° relative to the N state), whereas the head is captured in mid- and fully-swiveled conformation, (i.e. +8° and +16°, respectively) (Borovinskaya et al. 2008; Mohan et al. 2014). HygB binding moderately stabilized A-site tRNA binding and does not affect the formation of the A/P hybrid state (Peske et al. 2004). Previous reports indicate that addition of Par increases the fraction of ribosome in the N states (Wasserman et al., 2015). To estimate the kinetics of the reaction using ensemble kinetics, we mixed Kan or Par with PRE complexes in the absence of EF-G and observed in real time the SSU movements triggered by the antibiotics binding. For both drugs, we observed an increase of FRET efficiency indicating a stabilization of the SSU body and head in the N state (Figure 4A, B). Time courses with different Kan concentrations yielded a hyperbolic *k*_app_ dependence reaching a similar k_max_ for both SSU body and head movement, 160 ± 24 s^−1^ and 195 ± 13 s^−1^, respectively. The apparent affinity of the antibiotic to the PRE complex is about 30 ± 4 μM, and the lower limit for the association rate constant, k_a_ = 6 μM^−1^s^−1^ (see Methods). Ensemble kinetics did not show any signal change upon addition of HygB to the PRE complex, suggesting that at those conditions the conformational change is small (Supplemental Figure S1).

**Figure 4.**
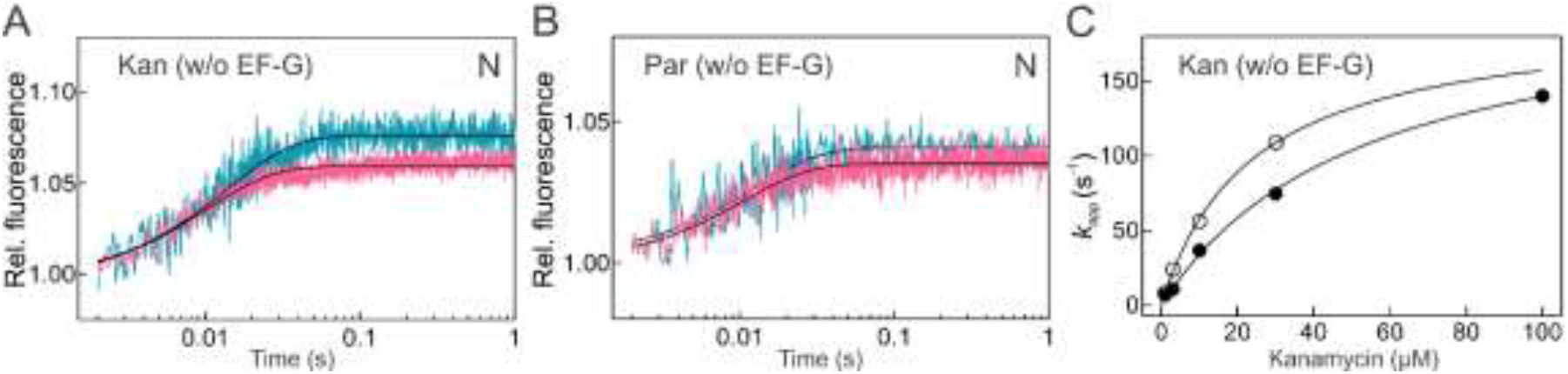
N state stabilization upon Kan or Par binding to PRE complex. Kan (A) and Par (B) effect on the PRE complex dynamics in the absence of EF-G. Body (teal) and head (pink) FRET fluorescence time courses were monitored in the stopped-flow apparatus upon addition of Kan (30 μM) or Par (5 μM) to PRE complexes. (C) Dependence of *k_apps_* values estimated by one-exponential fitting of time courses obtained with increasing concentrations of Kan (0.1 – 100 μM). Body (full black dots) and head (empty dots) Lines show hyperbolic fits. Error bars (which are smaller than symbol site) represent the error of the fit.

As with other aminoglycosides, binding of EF-G to PRE complexes in the presence of Par, Kan, or HygB does not impair GTP hydrolysis and Pi release (Peske et al. 2004). Kan and Par have a very similar effect on EF-G-facilitated motions of the ribosomal subunits (Figure 5A, B and Table 1). Because Kan and Par stabilize the N state, the amplitude of the forward SSU rotation upon EF-G binding is larger than without antibiotics; albeit at the cost of somewhat slower rates (Table 1). Once the ribosome completes the forward movements (N → R), the reverse motions (R → N and S → N) occur 15 to 30 times slower than in the absence of antibiotics. The R→N and S→N lifetimes with Kan are τ_body_ = 0.8 s^−1^and τ_head_ = 4.5 s^−1^, and with Par τ_body_ = 0.6 s^−1^and τ_head_ = 2.6 s^−1^. The lifetime of the CHI state also increases 20 and 10 times with Kan and Par, respectively. The reverse subunits motions in the presence of HygB are even slower: τ_body_ = 18 s^−1^and τ_head_ = 153 s^−1^, which increases the lifetime of the CHI state by more than 800-fold compared to the uninhibited reaction (Figure 5C and Table 1; *k*_app2body_ = 0.12 s^−1^, *k*_app3body_ = 0.034 s^−1^ and *k*_app2head_ = 0.015 s^−1^, *k*_app3head_ = 0.004 s^−1^). For all three drugs, the sequence of ribosome motions that lead to translocation is the same as in the absence of the drug (Figure 5). Additionally, the ratio of the apparent rate of reverse body to reverse head movement (i.e. *k*_app body_ / *k*_app head_) remains in the range between 5 and 10, which is similar for the unperturbed reaction. This suggests a certain degree of synchronization of the SSU reverse motions also when the antibiotics slow down these reactions.

**Figure 5.**
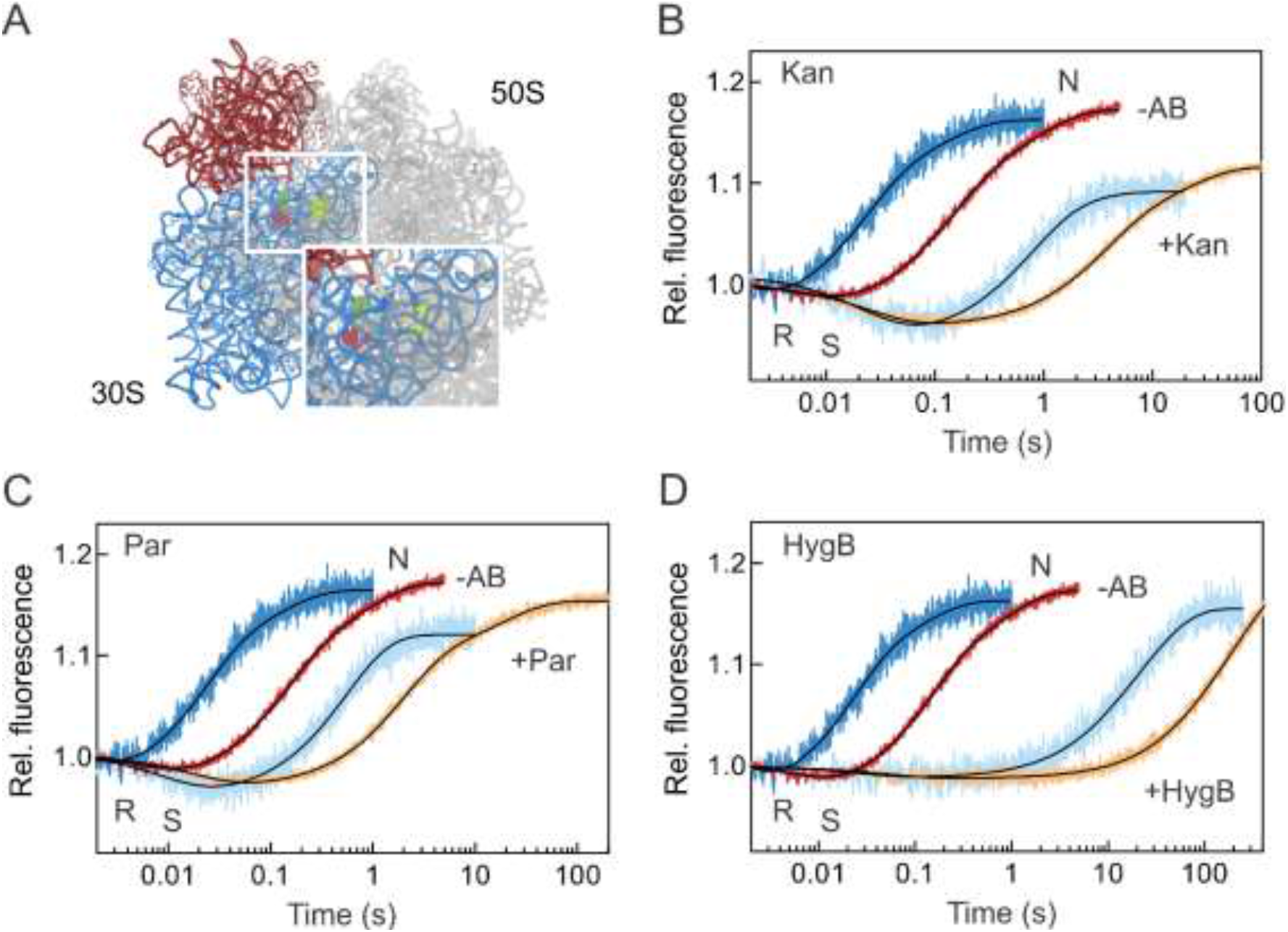
Kan, Par and HygB maintain the order of SSU body-head movements. (A) Par (salmon and yellow) and HygB (green) binding sites on the SSU. SSU body and head are highlighted in blue and red, respectively, the LSU is gray. EF-G-dependent SSU body (light blue) and head rotation (orange) were monitored in the presence of Kan (B), Par (C), or HygB (D) and compared to the reactions in the absence of antibiotics (−AB; blue is SSU body; red is SSU head).

### Neomycin

Neo is a 4, 5-aminoglycoside that can bind to multiple sites on the ribosome (Borovinskaya et al. 2007a; Wang et al. 2012; Zhou et al. 2014) (Figure 6A). smFRET studies suggest that in the absence of EF-G the effect of Neo is strongly concentration-dependent. At concentrations of Neo <2 μM, the antibiotic stabilizes PRE complexes in the N conformation, whereas at higher concentrations (>20 μM) the SSU body is trapped in an intermediate FRET state (Feldman et al. 2010; Wang et al. 2012).

**Figure 6.**
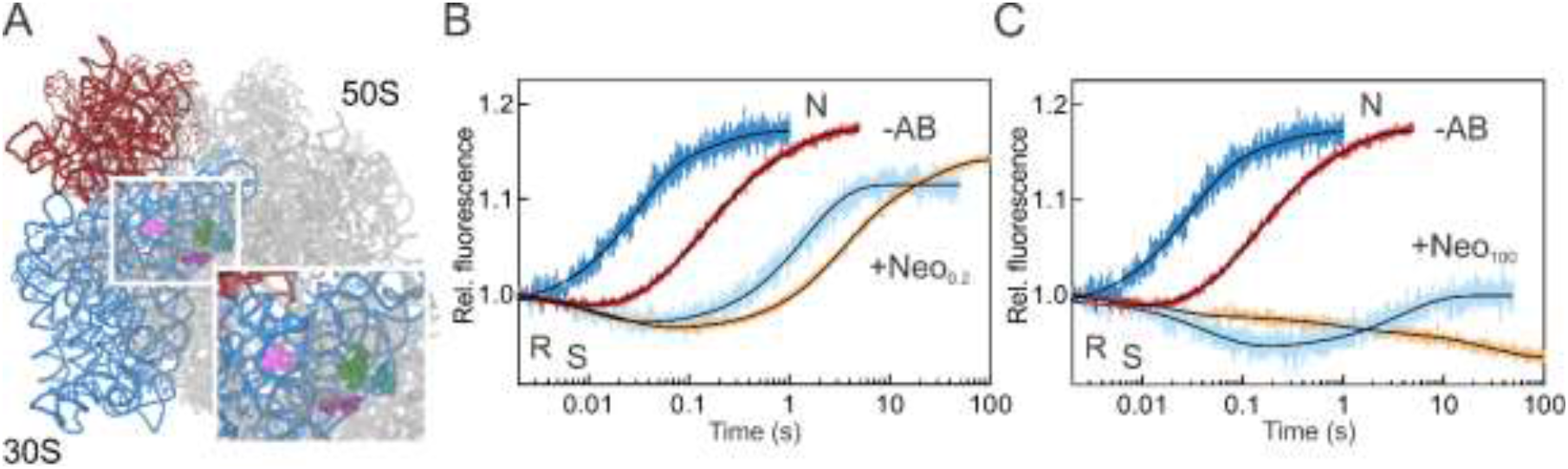
Neo concentration-dependent effects. (A) Neo binding sites on the SSU (pink and violet) and LSU (dark green). SSU body and head are highlighted in blue and red, respectively, the LSU is gray. EF-G-dependent SSU body (light blue) and head swiveling (orange) were monitored in the presence of 0.2 μM (B) or 100 μM (C) Neo and compared to the uninhibited reactions (−AB).

Structural and biochemical reports suggested that Neo does not inhibit EF-G binding to PRE complexes, but rather stabilizes the factor on the ribosome (Campuzano et al. 1979a; Campuzano et al. 1979b; Zhou et al. 2014). At low concentration, the effect of Neo is similar to that of Kan and Par. Upon EF-G binding, the sequence of the SSU body and head motions (N→R→N and N→S→N, respectively) does not change, but they are slower than in the absence of the drug (τ_body_ = 1.5 s^−1^and τ_head_ = 4.6 s^−1^, τ_CHI_ = 19 s^−1^; Figure 6B) (Wang et al. 2012). At higher Neo concentrations (100 μM), when also the LSU binding site is likely occupied by the drug, the early R state stabilization is still observed, however SSU head swiveling is extremely slow (Figure 6C). The reverse motions are almost completely abolished and the SSU head adopts an exaggerated S conformation similar to the translocation complexed trapped in an intermediate state of translocation in the presence of Neo and fusidic acid (Zhou et al. 2014).

### Streptomycin

Str binds at the decoding site of the ribosome, contacting h18, h27, and h44 of 16S rRNA as well as ribosomal protein S12 (Carter et al. 2000; Demirci et al. 2013) (Figure 7A). It increases the affinity of peptidyl-tRNA to the A site (Peske et al. 2004). Previous biochemical experiments suggested that Str induces a conformation of the PRE complex which is favorable for translocation; the conformational distortion caused by the opposite motions of the SSU body and head may account for the more open, translocation-prone conformation.

**Figure 7.**
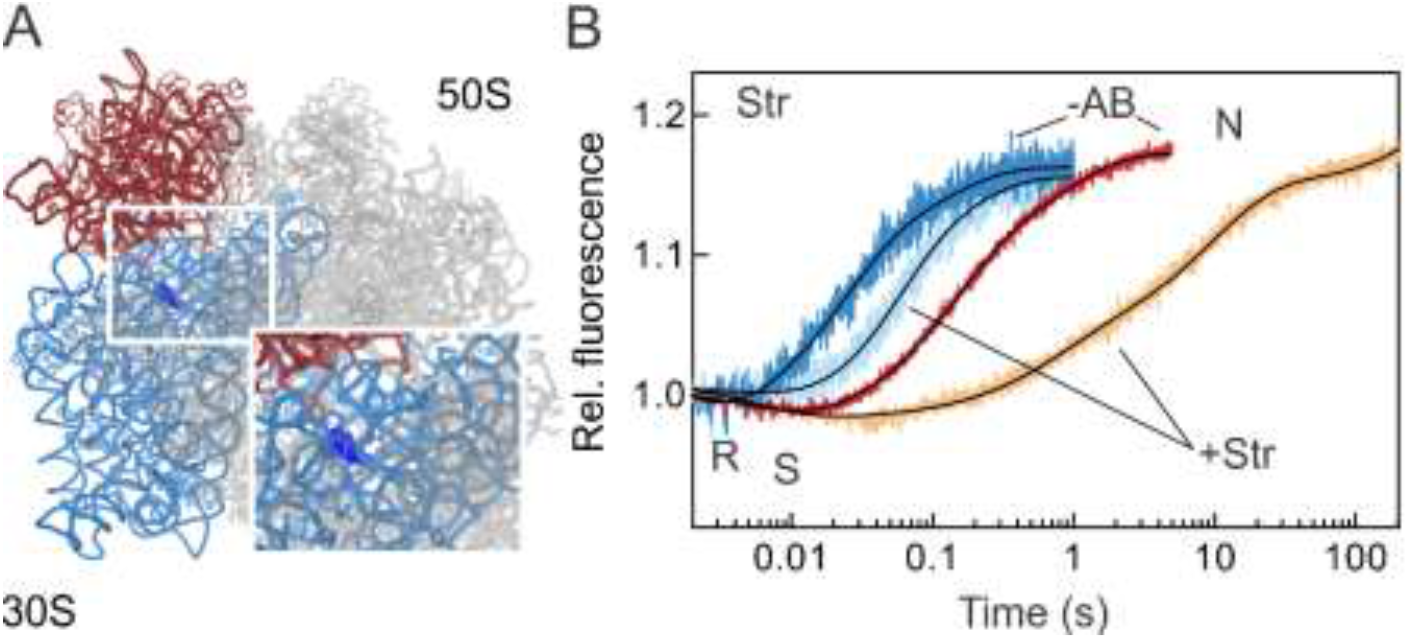
Str induces a long-lived translocation-prone state. (A) Str (dark blue) binding site on the SSU. (B) EF-G-dependent movements of the SSU body and head domains were monitored in the presence of Str (light blue and orange, respectively), and compared to those in the absence of the drug (−AB).

EF-G binding, GTP hydrolysis and Pi release are not affected by Str (Peske et al. 2004). The SSU body domain rotation upon addition of EF-G is only three-to-four times slower than the unperturbed reaction (Figure 7A and Table 1). The backward body rotation (R→N), which coincides with the formation of the CHI state, occurs with an apparent rate of 12 s^−1^ (τ = 70 ms). This result is in agreement with two previous studies where the rates of tRNA translocation on the LSU and SSU were estimated to be 7 and 11 s^−1^, respectively, under single round translocation conditions and in the presence of the inhibitor (Peske et al. 2004; Holtkamp et al. 2014). For the SSU head domain, the rate of the forward swiveling (manifested by the FRET decrease in Figure 5A) is unaltered by the presence of the drug. However, the back swiveling is very slow and appears multiphasic, with τ = 46 s, i.e. 240 times slower than in the absence of the drug (Table 1). Given that single round tRNA translocation in the presence of Str is much faster than that (Peske et al. 2004; Holtkamp et al. 2014), these data suggest that tRNAs move to their final destinations during the prolonged “unlocked” state of the ribosome. These data would, however, predict that multiple turnover of EF-G on the PRE complexes is affected, which in fact has been observed using the puromycin reaction (Peske et al. 2004). Thus, the small translocation defect caused by Str results from a combination of the A-site tRNA stabilization, formation of a translocation-prone conformation of the PRE complex, and the delay in the back SSU head domain swiveling.

## Discussion

This paper shows how antibiotics alter the movements of ribosomal subunits during translocation (Figure 8). SSU body domain rotation and head domain swiveling are key repetitive motions that promote the movement of the ribosome along the mRNA. Impairment of these motions eventually leads to translational arrest. Interestingly, although antibiotics binding can favor a particular orientation of ribosomal subunits, e.g. stabilize N or R or some intermediate state of the ribosome, none of those we tested seems to interfere with EF-G–GTP-induced forward SSU motions, regardless of the conformation favored in the absence of EF-G. This observation is in agreement with the finding that antibiotics do not impair EF-G binding and supports the notion that EF-G can bind to the ribosome in any conformation, and accelerate and stabilize the R conformation of the SSU. The antibiotics studied here have in common that they impair the backward movements of the SSU, but each of them affects the movements of the SSU body and head domains to a different extent. Vio blocks essentially all backward movements of the SSU, which in addition to its strong stabilizing effect on the A-site tRNA binding explains its strong inhibitory effect. Also Spc halts the ribosome before the reverse motions of the SSU body and head domains occur. However, the mechanism of action appears more complex with Spc, because it actually destabilizes A-site tRNA binding. Spc appears to disrupt the coupling between the GTPase and Pi release activity of EF-G and the tRNA-mRNA movement, altering the translocation pathway in ways similar to those in the absence of GTP hydrolysis. Neo, at high concentrations, essentially blocks SSU movements. In contrast, Kan, Par, HygB and Neo (at low concentration) do not block the movements completely, but make the reverse SSU body rotation and head swiveling much slower than during the uninhibited reaction. Importantly, the sequence of events, in which the reverse SSU body rotation precedes the reverse SSU head swiveling, is maintained, as backward SSU body rotation is 5 to 10 times faster than the backward SSU head swiveling. It thus seems that the coordination of the backward motions is still in play even when the reaction requires longer than 200 s instead of 1-2 s. One exception is Str, which has only a moderate effect of the SSU body rotation, but impairs the SSU head rotation. Notably, Str is not a strong translocation inhibitor, but could slow down the elongation cycle after tRNA translocation is completed. The present findings show how interfering with the principal subunit motions results in translocation inhibition.

**Figure 8.**
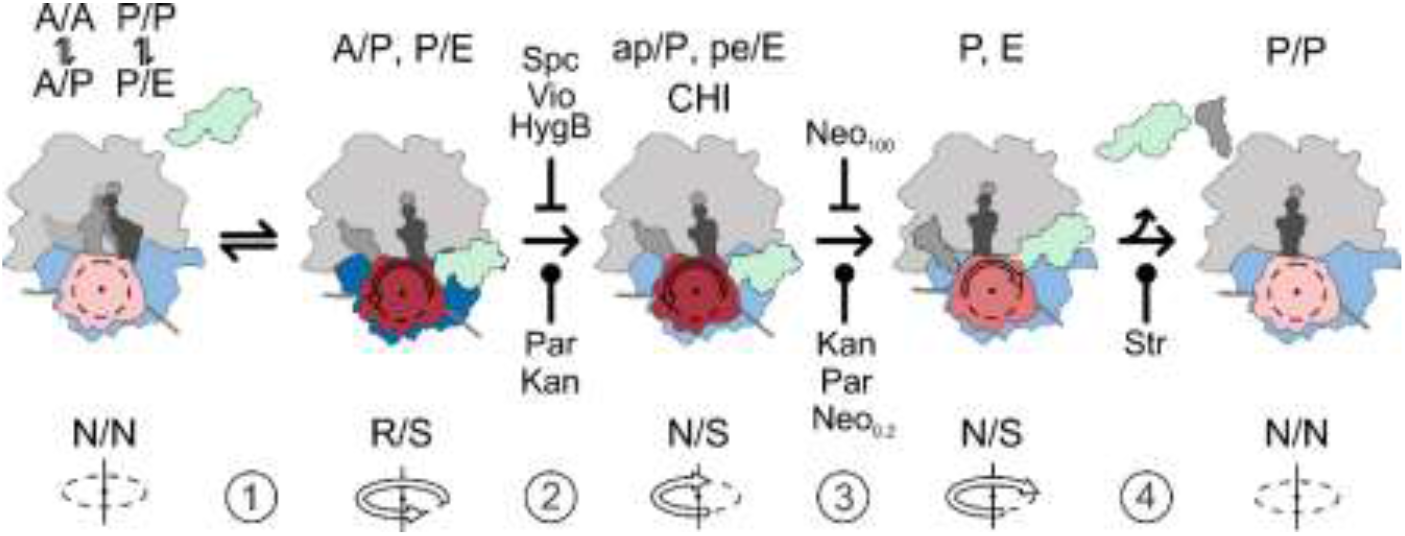
Summary of inhibition mechanisms of antibiotics impairing translocation. The inhibitory step for each antibiotic and the ability to halt (⊥) or slow down ( ) the motions of ribosomal subunits during translocation is indicated. We consider a step as inhibited if it is >300-fold slower than in the absence of antibiotics.

## Materials and Methods

### Ribosomes, mRNAs, tRNAs and translation factors

Ribosomal subunits were prepared from deletion strains (ΔS6, ΔS13, ΔL9, ΔL33) according to the protocol used for native ribosomes (Rodnina and Wintermeyer 1995). Initiation factors (IF1, IF2, IF3), EF-G, EF-Tu and tRNAs (f[^3^H]Met-tRNA^fMet^ and [^14^C]Phe-tRNA^Phe^) were prepared as described (Rodnina and Wintermeyer 1995; Savelsbergh et al. 2003; Cunha et al. 2013; Holtkamp et al. 2014). mRNA constructs were synthesized by IBA (Göttingen, Germany) using the sequence 5’- GUUAACAGGUAUACAUACUAUGUUUGUUAUUAC-3’.

### Labeling of ribosomal subunits

*E. coli* genes for proteins bS13, bL33 were PCR-amplified and cloned individually into the pET24(a) plasmid (Novagen). Similarly, *E. coli* genes for proteins bS6 and bL9 were PCR-amplified and cloned individually into the pET28(a) plasmid (Novagen). Recombinant single-cysteine proteins were then expressed, purified, labeled and refolded as previously described (Cunha et al. 2013; Belardinelli et al. 2016a; Sharma et al. 2016). Reconstitution of labeled subunits was carried out as described (Ermolenko et al. 2007a; Ermolenko et al. 2007b; Belardinelli et al. 2016a; Sharma et al. 2016). Briefly, purified ΔS13, ΔS6, ΔL9 and ΔL33 ribosomal subunits were reconstituted with a 1.5-fold excess of labeled S13, S6, L9 and L33, respectively. The excess of labeled protein was removed by ultra-centrifugation through a 30% sucrose cushion in TAKM_7_ buffer (50 mM Tris-HCl pH 7.5, 70 mM NH_4_Cl, 30 mM KCl, 7 mM MgCl_2_. The extent of ribosomal subunit labeling, as determined spectrophotometrically, was about 95-100%.

### Preparation of the complexes

Preparation and purification of PRE complexes were carried out as described (Rodnina et al. 1997; Cunha et al. 2013; Holtkamp et al. 2014; Belardinelli et al. 2016a; Sharma et al. 2016). Briefly, labeled SSU were first heat activated in TAKM_21_ buffer (i.e. with 21 mM MgCl_2_) for 30 min at 37°C. Activated SSU were then mixed with a 1.5-fold excess of labeled LSU, a 3-fold excess of mRNA, a 2-fold excess of IF1, IF2, IF3 each, and a 2.5-fold excess of f[^3^H]Met-tRNA^fMet^ in TAKM_7_ buffer containing 1 mM GTP, and incubated for 30 min at 37°C. Ternary complex EF-Tu–GTP–Phe-tRNA^Phe^ was prepared by incubating EF-Tu (2-fold excess over tRNA) with 1 mM GTP, 3 mM phosphoenolpyruvate and 0.1 mg·ml^−1^ pyruvate kinase for 15 min at 37°C prior to the addition of aa-tRNA. PRE complexes were assembled by mixing initiation complexes with a 2-fold excess of ternary complex and incubated for 1 min at 37°C. The resulting PRE complexes were purified by ultra-centrifugation through 1.1 M sucrose cushion in TAKM_21_. Pellets were resuspended in the same buffer and the efficiency of PRE complex formation was between 70 and 80% as estimated by nitrocellulose filtration and radioactivity counting.

### Rapid kinetics

Rapid kinetics experiments were carried out using a stopped-flow apparatus (SX-20MV; Applied Photophysics) in TAKM_7_ at 37°C. To monitor SSU head swiveling and body rotation, we used double-labeled ribosomes (S13Atto540Q–L33Alx488) and (S6Alx488–L9Alx568), respectively (Belardinelli et al. 2016a; Sharma et al. 2016). Alx488 fluorophores were excited at 465 nm and the emission was recorded after passing through a KV500 cut-off filter (Schott). When FRET between Alx488 and Alx568 was monitored, the emission of the acceptor fluorophore was recorded after passing a KV590 filter. To study the effect of antibiotics on SSU rotation in the absence of EF-G, dually-labeled PRE complexes (0.05 μM) were rapidly mixed with the following concentrations of antibiotics (HygB, 20 μM; Str 20 μM; Spc 1 mM; Kan 100 μM; Par, 5 μM; Neo 0.2 and 100 μM, and Vio 200 μM). SSU rotation upon EF-G–induced translocation was monitored after mixing PRE complexes (0.05 μM) with saturating concentration of EF-G (4 μM) and GTP (1 mM), previously pre-incubated with the respective antibiotics (see above). The concentration dependencies of SSU head swiveling and body rotation were assessed upon mixing PRE complex (0.05 μM) with increasing concentrations of Kan (1, 3, 10, 30 and 100 μM). Time courses were evaluated by one-, two- or three-exponential fitting, as appropriate, using GraphPad Prism software. Weighted average was calculates as τ = (*A*_2_ + *A*_3_)/(*k_app2_* × *A*_2_ + *k_app3_* × *A*_3_). S.e.m. associated to the kinetics were calculated by the GraphPad Prism software. Each time course reflects the average of 7 to 10 technical replicates.

## Acknowledgements

We thank Wolfgang Wintermeyer for critical reading the manuscript; Rachel Green and Janine Maddock for the *E. coli* strains carrying ΔS13 and ΔL33 ribosomes, respectively; Anna Pfeifer, Olaf Geintzer, Sandra Kappler, Christina Kothe, Theresia Steiger, Franziska Hummel, Tessa Hübner, Vanessa Herold and Michael Zimmermann for expert technical assistance. Funding: The work was funded by the Max Planck Society and the Deutsche Forschungsgemeinschaft in the framework of Sonderforschungsbereich 860 (M.V.R.). H.S. acknowledges a Boehringer Ingelheim Doctoral Fellowship.

---

**Supplemental Figure S1.**
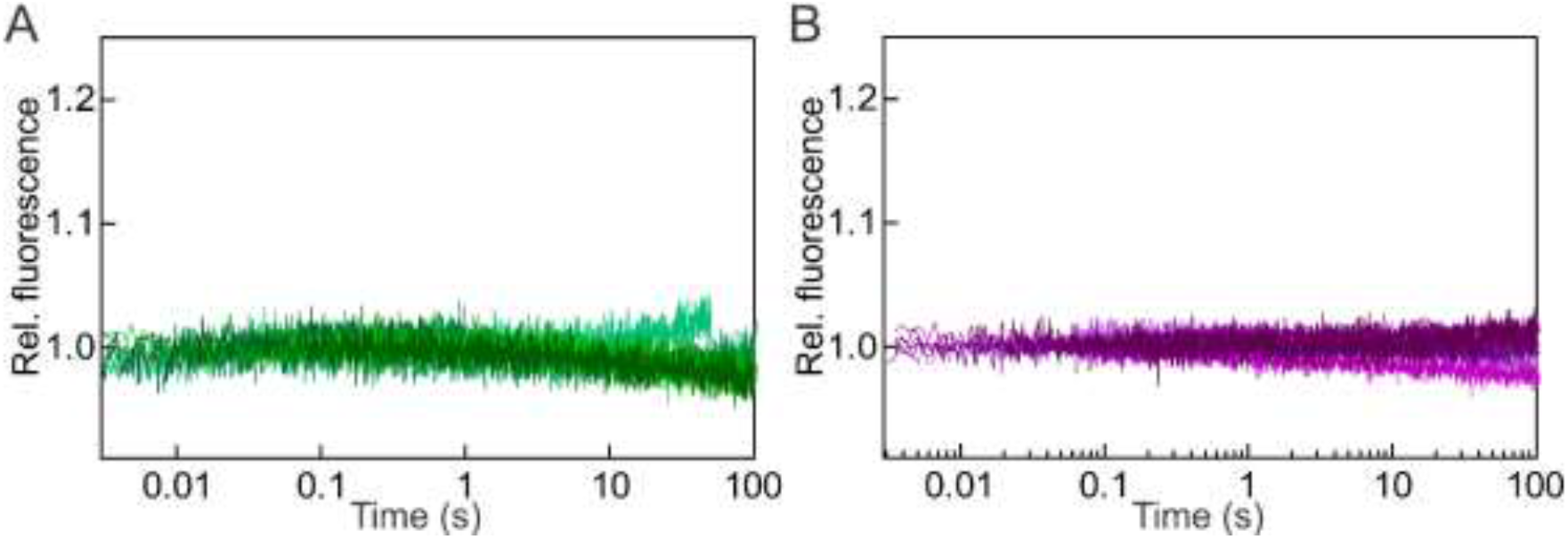
Body (A; green shade) and head (B; purple shade) control traces of PRE complex mixed with either buffer or antibiotics. Darkest traces are PRE complex mixed with buffer, then towards the lightest shade of either green (body) or purple (head) are the control traces with Spc, Str, HygB, Vio & Neo. (Par & Kan are omitted because they are shown in Fig. 4).

